# Mobile Type VI secretion system loci of the gut Bacteroidales display extensive intra-ecosystem transfer, multi-species sweeps and geographical clustering

**DOI:** 10.1101/2021.01.21.427628

**Authors:** Leonor García-Bayona, Michael J. Coyne, Laurie E. Comstock

## Abstract

The human gut microbiota is a dense microbial ecosystem with extensive opportunities for bacterial contact-dependent processes such as conjugation and type VI secretion system (T6SS)-dependent antagonism. In the gut Bacteroidales, two distinct genetic architectures of T6SS loci, GA1 and GA2, are contained on integrative and conjugative elements (ICE). Despite intense interest in the T6SSs of the gut Bacteroidales, there is only a superficial understanding of their evolutionary patterns, and of their dissemination among Bacteroidales species in human gut communities. Here, we combine extensive genomic and metagenomic analyses to better understand their ecological and evolutionary dynamics. We identify new genetic subtypes, document extensive intrapersonal transfer of these ICE to Bacteroidales species within human gut microbiomes, and most importantly, reveal frequent population sweeps of these newly armed strains in multiple species within a person. We further show the distribution of each of the distinct T6SSs in human populations and show there is geographical clustering. We reveal that the GA1 T6SS ICE integrates at a minimal recombination site leading to their integration throughout genomes and their frequent interruption of genes, whereas the GA2 T6SS ICE integrate at one of three different tRNA genes. The exclusion of concurrent GA1 and GA2 T6SSs in individual strains is associated with intact T6SS loci and with an ICE-encoded gene. By performing a comprehensive analysis of mobile genetic elements (MGE) in co-resident Bacteroidales species in numerous human gut communities, we identify 177 MGE that sweep through multiple Bacteroidales species within individual gut microbiomes. We further show that only eight MGE demonstrate multi-species population sweeps in as many human gut microbiomes as the GA1 and GA2 ICE. These data underscore the ubiquity and rapid dissemination of mobile T6SS loci within Bacteroidales communities and across human populations.

## Introduction

The order Bacteroidales encompasses numerous genera including the *Bacteroides, Parabacteroides* and *Prevotella*, which collectively are the most abundant Gram negative bacteria of the healthy colonic microbiota of many human populations. These bacteria secrete anti-bacterial proteins that antagonize closely related strains and species, providing a competitive advantage in the gut ecosystem (reviewed (Garcia-Bayona and Comstock, 2018)). Type VI secretion systems (T6SSs) are an antagonistic system of these gut bacteria. T6SSs are contractile nanomachines that inject toxic effectors into bacterial or eukaryotic cells. There are three distinct genetic architectures of T6SS in gut Bacteroidales termed genetic architecture 1, 2 and 3 (GA1, GA2, and GA3). GA3 T6SS loci are found exclusively and at high proportion in *B. fragilis* strains, and contain by two variable regions containing genes encoding effector and immunity proteins (Coyne, 2016). The effectors of distinct GA3 T6SS have potent killing activity (Chatzidaki-Livanis et al., 2016; Hecht et al., 2016; Wexler et al., 2016), targeting nearly all gut Bacteroidales species analyzed (Chatzidaki-Livanis et al., 2016). GA3 T6SSs were shown to be enriched among strains colonizing the infant gut and associated with increased abundance of *Bacteroides* in the human gut microbiota (Verster et al., 2017).

The GA1 and GA2 T6SS loci are contained on Integrative and Conjugative Elements (ICE) and are present in diverse Bacteroidales species (Coyne, 2016). In Proteobacterial species, T6SS loci are typically contained on non-core genomic islands, but, with few exceptions (Marasini et al., 2020), are rarely found on conjugative elements. Some proteobacterial T6SS-associated genes, such as immunity genes (Ross et al., 2019), have been identified on mobile elements and a full T6SS locus is present on a mobile prophage-like element in environmental *Vibrio cholera* strains (Santoriello et al., 2020). The presence of complete Bacteroidales T6SS loci on ICE allows for their distribution to other co-resident Bacteroidales species in the human gut.

We previously identified 48 human gut Bacteroidales strains of 13 different species that contain GA1 T6SSs. The ICE containing the GA1 T6SS loci are approximately 95% identical at the DNA level between different strains. We previously showed that GA1-containing ICE were transferred to several Bacteroidales species in the gut of two human subjects, respectively (Coyne, 2016; Coyne et al., 2014). Like the GA3 T6SSs, the GA1 T6SS loci contain two variable regions that encode identifiable effector and immunity proteins (Coyne, 2016). To date, the target cells antagonized by the GA1 T6SSs have not been conclusively identified.

The GA2-containing ICE are distinct from the ICE containing GA1 T6SS loci. ICEs containing GA2 T6SS are less identical to each other than are the GA1 ICE. The GA2 T6SSs contain three variable regions with genes encoding potential effector and immunity proteins. Unlike the GA1 T6SS loci, we did not previously detect the same GA2 T6SS in multiple species of an individual, and therefore had no evidence of its transfer to co-resident species within the human gut microbiota. Although *B. fragilis* strains can harbor both a GA1 and a GA3 T6SS locus, we did not identify a Bacteroidales strain that harbors a GA2 T6SS locus along with either a GA1 or GA3 T6SS locus, suggesting exclusion.

The present study was designed to address numerous important and outstanding questions regarding the T6SS of the gut Bacteroidales by in-depth analyses of genomic and metagenomic data. In this study, we show five sub-types of GA2 T6SS, show that GA2 T6SS loci are globally dominant and transfer to multiple species within a gut microbiota with population sweeps. We identify the insertion sites of both GA1 and the different GA2 subtype ICE and find their exclusion is associated with both T6SS genes and a gene encoding a protein with both *N*^6^-adenine methylase and SNF2 helicase domains. In addition, we identify 177 mobile genetic elements that undergo multi-species sweeps, but few to the extent of the GA1 and GA2 ICE that commonly sweep Bacteroidales communities in the human gut of diverse populations.

## Results and Discussion

### Identification of five different GA2 subtypes

Unlike the GA1 and GA3 T6SS loci that are highly identical (∼95%) outside of the effector and immunity gene regions, the GA2 T6SS loci are more variable with less than 80% DNA identity between some loci (Coyne, 2016). To better study this variability, we compared the conserved regions (excluding the effector and immunity genes, Fig 1A) of 45 different GA2 loci and found that they segregate into 5 distinct subtypes (GA2a-e). With the exception of GA2e, DNA-level identities within a subtype are high (>97%), whereas cross-type identities generally run in the low 80% (subtypes GA2b and GA2c are more alike, demonstrating ∼89% identity) (Fig 1B). Each GA2 subtype clearly segregates to a distinct branch of a phylogenetic tree (Fig 1C), yet each retains the gene order and functions characteristic of the GA2 architecture, further supporting this subtyping.

**Figure 1.**
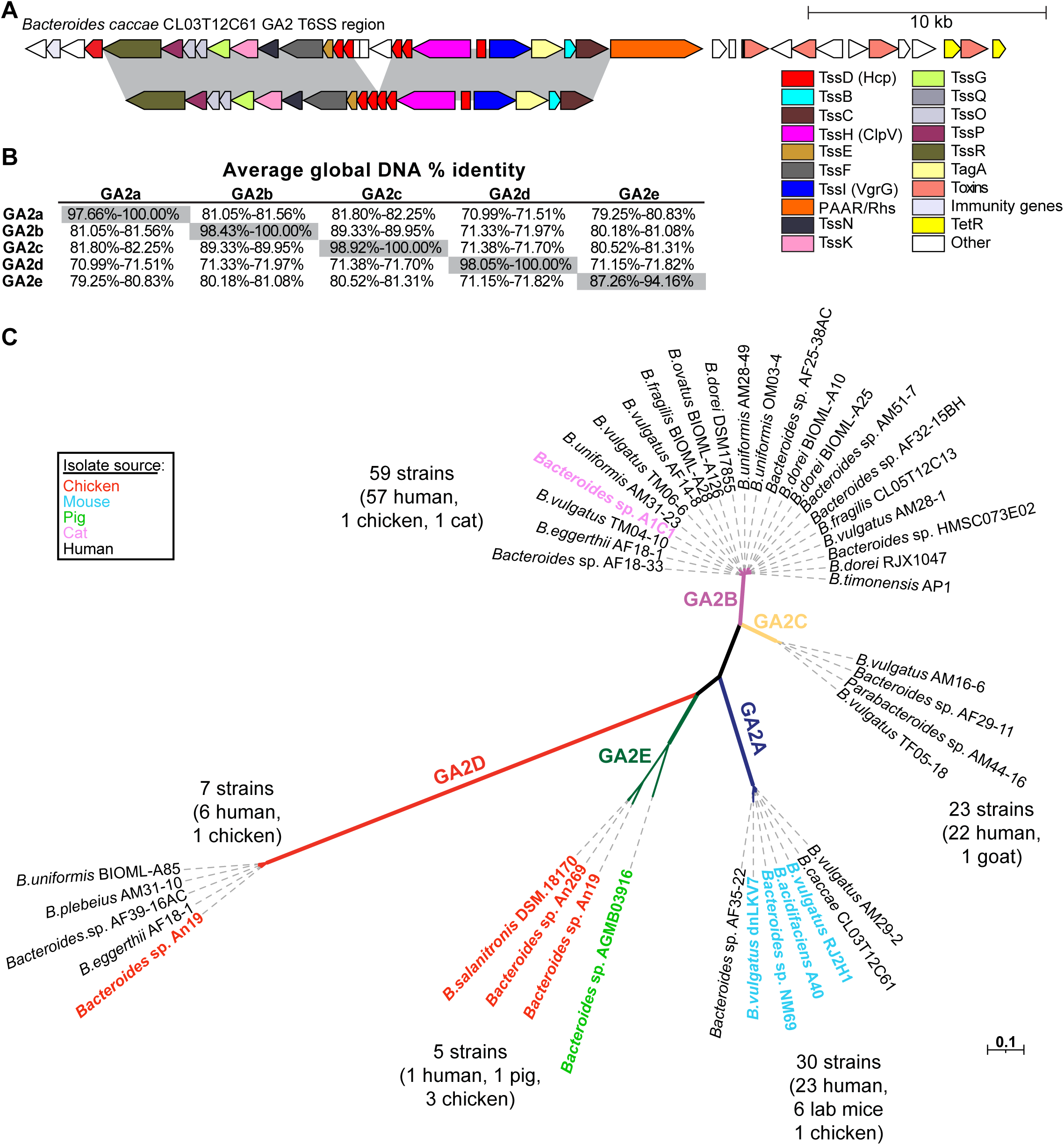
GA2 segregate into five different subtypes. **A**. Schematic of the GA2 genetic architecture of *B. caccae* CL03T12C61 as example. Segments highlighted in gray represent the conserved regions made into a concatemer used for the analyses shown in panels B and C. **B**. Average global DNA percent identity between the conserved concatemers of different GA2 subtypes. **C**. Maximum likelihood phylogenetic tree of the GA2 regions showing the separation into 5 subtypes. Only some strains are shown with the numbers at the end of each branch the total number of isolate genomes, out of the 1434 Bacteroidales genomes queried, that contain that subtype of GA2. Some isolates from non-human sources are highlighted in colored font.

### Prevalence of GA1, GA2, and GA3 T6SS loci in sequenced Bacteroidales isolates from human, animal, and environmental sources

Our initial report of the prevalence of the three distinct genetic architectures of T6SSs in human gut Bacteroidales was performed with 205 human gut Bacteroidales genome sequences that were available in 2015 (Coyne, 2016). To provide a more comprehensive analysis, we include here genome sequences of all Bacteroidales strains from any source available as of June, 2020. We attempted, with the information available to us, to include only genomes from isolated bacteria excluding those assembled from metagenomic sequences, and to reduce redundant genomes (multiple longitudinal isolates of the same strain from the same person or other identical strains sequenced multiple times). The final set includes 1434 Bacteroidales genomes of 14 different families and 41 different genera (Table S1). These genomes were queried with concatemers of GA1, GA2a, GA2b, GA2c, GA2d, GA2e, and GA3 T6SS loci with the divergent genes removed (Fig S1). Of these 41 genera, we detected GA1, GA2, and GA3 T6SSs in only three genera, *Bacteroides, Parabacteroides* and *Prevotella*, and nearly all of these strains were from gut sources (Table S1). As previously described, and reinforced by this analysis, the GA3 T6SSs are present exclusively in *B. fragilis*, with 93 of the 126 *B. fragilis* genomes analyzed here containing a GA3 T6SS locus. The GA1 T6SSs loci were found to be relatively evenly distributed among *Bacteroides* and *Parabacteroides* genomes, with no obvious species preference (Fig. 2); however, none of the 26 *Prevotella copri* genomes contains a GA1 locus, whereas both of the sequenced *P. stercorea* genomes do. In our previous analysis of 205 genomes, only nine genomes were found to contain a GA2 T6SS loci, suggesting then that GA2 loci may be rare in gut Bacteroidales. This expanded analysis shows that there are nearly as many GA2 loci (124) as GA1 loci (129) in these sequenced strains. Unlike the GA1 loci, this analysis reveals a GA2 species-level distribution bias, with GA2 loci being found at high prevalence in some species; *B. eggerthii* (43% of strains), *B. vulgatus* (40%), *B. uniformis* (30*%), B. stercoris* (23%), and *B. dorei* (26%); and more rarely in other species; *B. ovatus* (2%), *B. fragilis* (1.6%), *B. thetaiotaomicron* (0%), *B. intestinalis* (0%) (Fig. 2, Table 1). Among the GA2 subtypes, GA2b is the most prevent in this genome set, present in 62 genomes, followed by GA2a (26 genomes), GA2c (24 genomes), GA2d (7 genomes), and GA2e (5 genomes).

**Figure 2.**
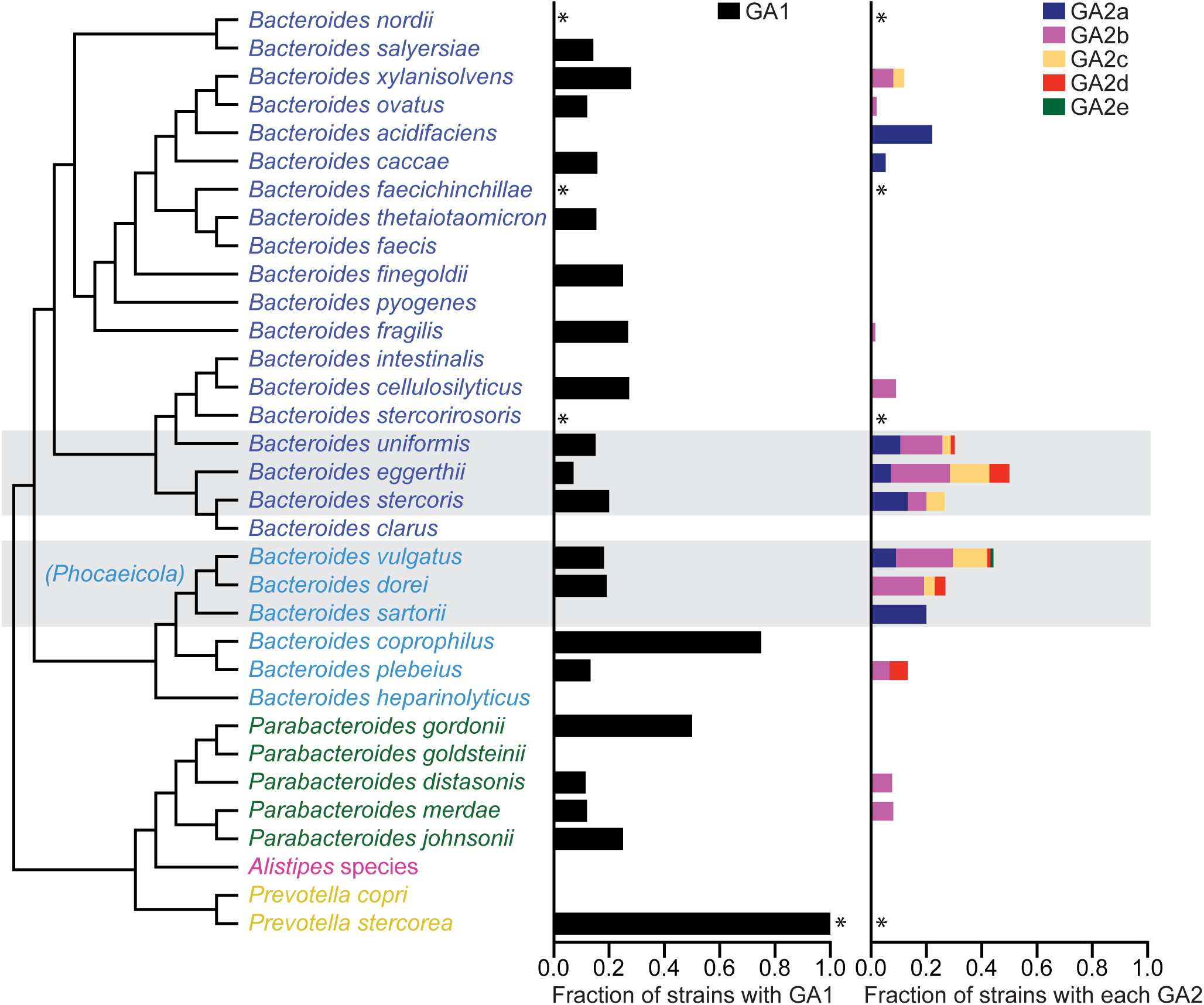
Abundance of GA1 and GA2 loci in different Bacteroidales species. Shown is the fraction of strains within each species harboring each type of GA. *Three or less sequenced isolates. Strains with no species call were not included.

The absence of GA1 and GA2 loci in non-gut Bacteroidales species, such as those that occupy the oral cavity or vagina, could be due to lack of cell-cell contacts, but both oral and vaginal Bacteroidales can occupy the gut microbiota, and even at low frequency, transfers to Bacteroidales in other ecosystems would be expected to happen. This observation suggests a fitness advantage conferred by the T6SSs that is unique to the gut. An interesting finding is the presence of some T6SS loci in the genomes of non-human isolates. Of the five GA2e loci detected, only one is from a human isolate, three are present in isolates from the ceca of chicken, and one from the stool of a pig (Table S1). One of the seven GA2d loci is from a strain isolated from a chicken. Of the 26 isolates with GA2a loci, six are mouse isolates, and one of the 62 strains with GA2b loci was isolated from a cat, and one from a chicken. The data suggest that the GA2e subtype is of non-human origin as only one human isolate contained a GA2e locus. In contrast, the few non-human strains containing the other four GA2 subtypes may have been acquired by these animals from a human source, as they were isolated from domestic or lab animals.

### Intra-ecosystem GA2 ICE transfers to co-resident species

In our previous analyses of the Bacteroidales strains from four healthy human volunteers, we showed that a GA1 ICE transferred between multiple *Bacteroides* and *Parabacteroides* species in the gut of two of these individuals. In contrast, we observed no intra-community transfer events for GA2 ICEs (Coyne, 2016). To determine if we could identify GA2 ICE transfers among co-resident species in the human gut, we screened the *Bacteroides* and *Parabacteroides* strains of four additional healthy human volunteers previously isolated as part of a longitudinal study (Zitomersky et al., 2011b). Using PCR primers that amplify a 675-bp conserved region of GA2 loci (Fig. 3A), we detected GA2 regions in numerous isolates from three of the four communities analyzed: CL06, CL08, and CL11 (Fig. 3B). We PCR-amplified a ∼2.7 kb variable region of these GA2-positive strains (Fig. 3A) and sequenced the amplicons. These DNA regions were identical between species of the same community, but differed between the three communities.

**Figure 3.**
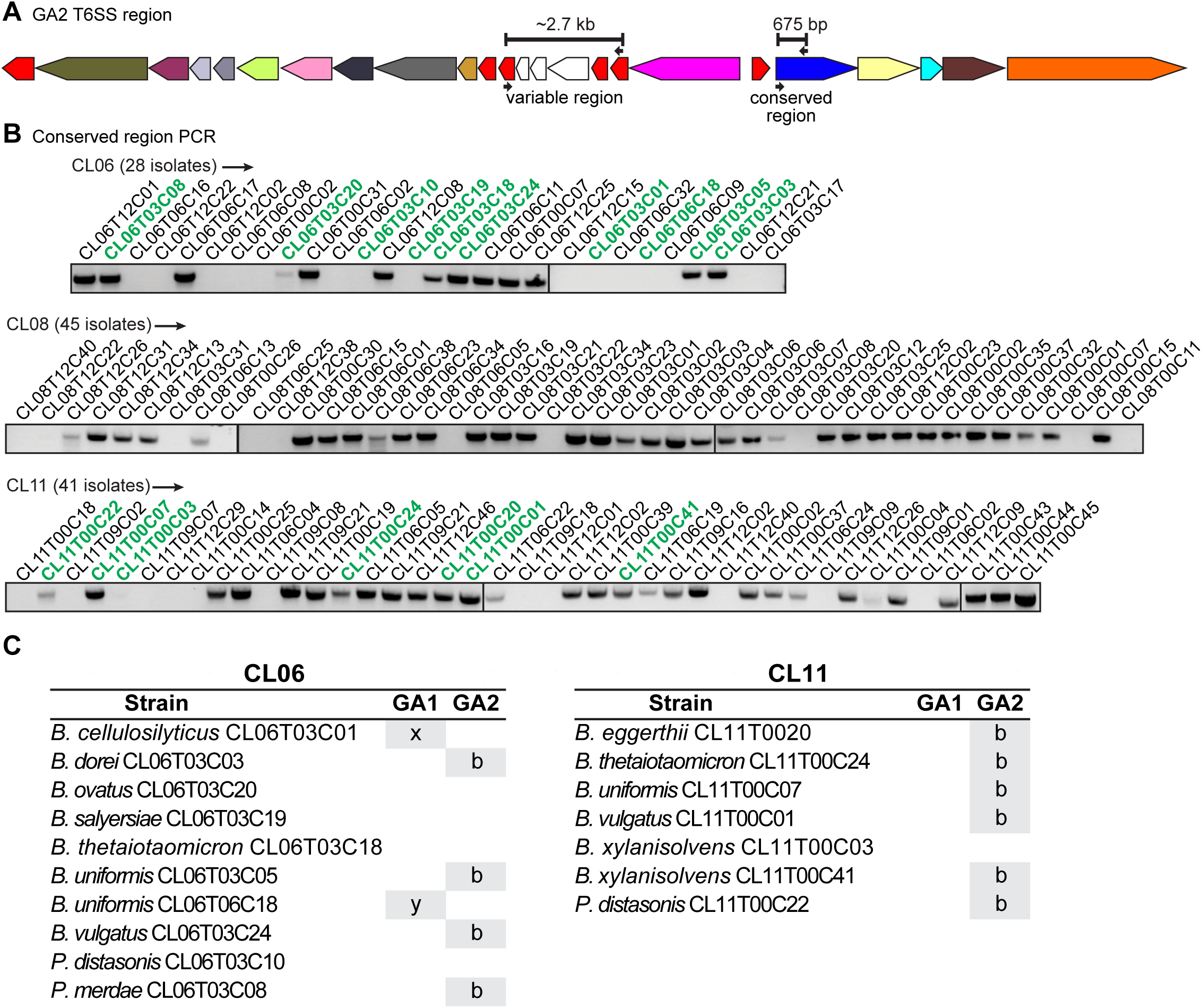
Screen for intra-community GA2 transfer events in the Comstock lab strain collection. **A**. Schematic of a GA2 T6SS. Small arrows indicate screening primer annealing sites and bars indicate the corresponding PCR product. Color scheme as described in Fig 1. **B**. PCR screen of the isolates from the three communities indicated, using primers for the GA2 conserved region. Strains bolded in green were selected for whole genome sequencing. **C**. Strains sequenced in this study and whether they contain GA1 or GA2 T6SS loci. x and y in GA1 column indicate the regions are distinct.

Using the previously determined species designation of each isolate (Zitomersky et al., 2011b), we selected strains representing all species from communities CL06 and CL11 and sequenced their genomes using SMRT sequencing (Fig. 3C). Sequence analyses showed 99.99-100% DNA identity of the T6SS and ICE regions among all six GA2+ species of ecosystem CL11 (out of seven total species) and determined that they are of the GA2b subtype. The GA2b region is intact in all strains except for *B. uniformis* CL11T00C24, in which a transposase insertion interrupted the first Rhs toxin gene. Of note is the identification of a *B. thetaiotaomicron* strain (BtCL11T00C24) containing a GA2 locus: this is the first *B. thetaiotaomicron* strain identified with a GA2 locus. Of the four CL06 species containing a GA2, all loci are 100% identical, except for a 445 bp duplication in the *tssC* gene in three strains but absent in *B. uniformis* CL06T03C05. These data clearly demonstrate GA2 ICE transfer within the human gut, with subsequent community sweeps. For the CL06 community, two species lack the GA2 locus, but each has a distinct GA1 T6SS loci, demonstrating lack of community transfer of these GA1 ICE.

To further study GA1 and GA2 ICE sweeps in a larger set of communities, we analyzed three available datasets of sequenced gut bacterial isolates from healthy adults: BIOML (longitudinal study from USA) (Poyet et al., 2019), CGR (China) (Zou et al., 2019) and UK (Browne et al., 2016). From these datasets, we analyzed the Bacteroidales isolates from subjects from whom at least three different Bacteroidales species were isolated, yielding 10 subjects from the BIOML set, 12 from the UK set, and 71 from the CGR set (Fig. 4A, Table S3). We searched each genome collection for the GA1 T6SS region and for each of the GA2 T6SS subtype regions using the same concatemer queries and blastn parameters as were used during analysis of the NCBI genome set. We assumed a recent (i.e. within the lifetime of the participant) intra-host transfer event if the regions were >99.95% identical at the DNA level between two species and a multi-species sweep if the identical region was found in at least three different species from the same community. For GA1, intra-community transfer limited to two species was observed in nine CGR and two BIOML communities (Fig 4, Table S3, S4). For GA2, intra-community transfer limited to two species was observed in two BIOML, one UK and eight CGR communities. Interestingly, for subject AF15 from the CGR set, two strains (*B. uniformis* AF15-14LB and *B. vulgatu*s AF15-6A) each contain both a GA2b and a GA2c, integrated at different chromosomal sites. This same phenomenon is observed in individual AF16, where both *B. uniformis* AF16-7 and *B. vulgatus* AF16-11 have very high sequence similarity across the whole genome with their AF15 cognates. These data indicate inter-person transfer of bacteria and that the GA2 transfer events occurred prior to passage of these bacteria to the recipient.

**Figure 4.**
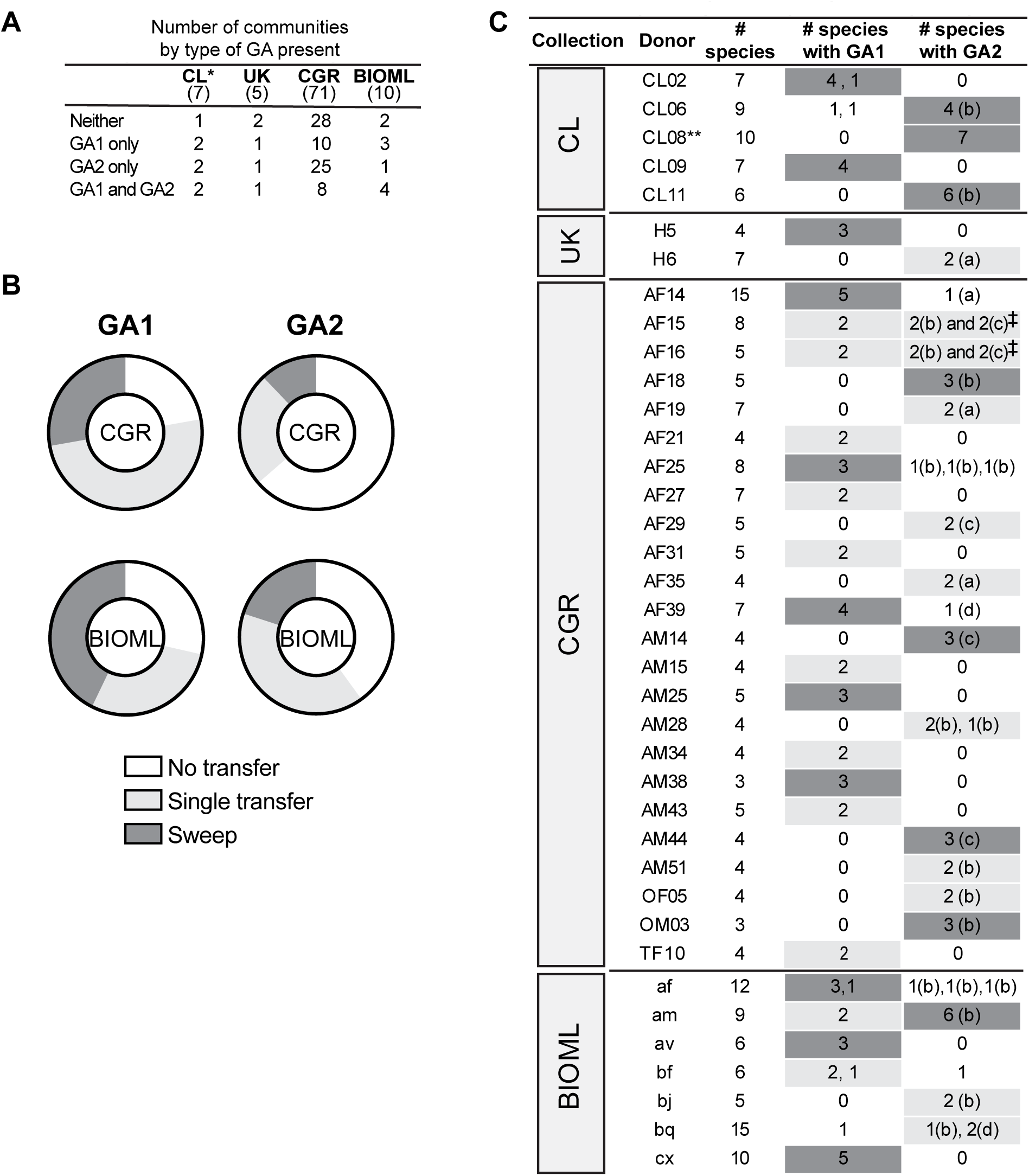
Intra-community sweeps are common for GA1 and GA2. **A**. Number of communities per dataset, classified by GA1 and GA2 presence/absence. Numbers in parentheses under each dataset name indicate total number of volunteers analyzed in that collection. *Includes CL08 and CL14 which were screened by PCR as described in the text. **B**. Percentage of communities in the CGR and BIOML collections that show the indicated event (no transfer, single transfer, multi-species sweep), out of the total communities in that dataset that have at least one strain with the indicated GA type. **C**. Communities with a single intra-community transfer event (highlighted in light grey) or a multi-species sweep (highlighted in dark grey). Letters in parentheses indicate type of GA2. Numbers listed separately in an individual box indicate the regions are distinct loci. **\*\***Screened by PCR and sequencing of variable region, as described in the text. Species determined by 16s rRNA sequence. ‡The two relevant strains from donor AF15 and AF16 are nearly clonal and appear to have both a GA2b and GA2c (see details in text).

We determined that GA1 and GA2 multi-species sweeps (three or more species) are common (Fig. 4). For communities containing a strain with a GA1 locus, multi-species sweeps occurred in 27.8% of the CGR and 42.9% of the BIOML communities. For communities containing a species with a GA2 locus, 12.1% of the CGR and 20% of the BIOML communities showed evidence of sweeps, whereas no multi-species sweeps where observed in the UK communities. Simultaneous GA1 and GA2 sweeps were never observed suggesting some degree of exclusion. GA1 may be more readily transferred and fixed in a bacterial population than GA2, since for all datasets, the majority of communities with a GA1 ICE transferred it to at least one other species (Fig. 4B, C). In contrast, a majority of communities from the CGR dataset with a GA2 locus did not show evidence of transfer events. This phenomenon could be partially attributable to the species that are present in a particular community, as GA2 demonstrates a species bias. However, this does not fully account for the observed variation, as species that frequently contain GA2 are present in many of the communities with no transfer. It is therefore possible that transfer and fixation in the population may be limited by specific strain physiology and other ecosystem factors.

The longitudinal BIOML dataset comprises many isolates (often more than 20) of the same species per individual community. In most cases these isolates are nearly isogenic (designated here as the same strain), but sometimes two or more distinct strains of the same species coexist at the same collection time-point (Poyet et al., 2019). This allowed us to evaluate the completeness of sweeps within the population of a strain. GA1 sweeps were complete within a strain (all closely-related isolates had the GA1), with a conserved insertion site, indicating they originated from a single transfer event (discussed in the following section). We observed only one exception, where near-isogenic isolates with and without a GA1 ICE were detected (Table S5A, S5C, partial sweep bolded in S5C). Interestingly, two or more unrelated co-existing strains of the same species often had differences in GA1 presence or absence (Table S5C). That is to say, we observed five instances in which all isolates of one strain within a community had a GA1 while all the isolates of the other co-resident species-matched strain did not. The acquisition of a GA2 ICE did not always lead to a full sweep in the population of that strain, as we identified three instances of near isogenic co-resident strains some with and some without the GA2 ICE (two for a GA2b and one for a GA2d) (Table S5A, S5B). Similar to GA1, we also detected four instances of communities harboring two different co-existing strains of the same species, one with and one without the GA2 ICE. Finally, we identified four examples of GA2 within-species sweeps, and one GA1 within-species sweep, where some of the isolates of one strain lost a large part of the T6SS region, while in others, it remained intact. All together, these results show that both T6SS genetic architectures contained on ICEs often transfer horizontally in the gut and are fixed in the population of multiple species. We were unable to detect events were a species or strain previously devoid of a T6SS acquired one during the sampling time, perhaps due to the short sampling time of 1.5 years or less. Individual transfer events may be infrequent or may occur early upon strain acquisition or during population bottlenecks.

Acquisition of an ICE carries with it the cost of maintaining a large genetic element, but the widespread nature of ICE community sweeps suggests that the fitness benefit of its acquisition outweighs the metabolic cost. Despite numerous examples of ICE transfers and multi-species sweeps, there are also examples where no transfer or single transfers occurred, especially for the GA2 ICE. Therefore, fitness conferred by T6SS ICE acquisition is likely context-dependent and may be contingent upon host physiology and emergent properties of the community and its microbial composition.

### GA1 and GA2 ICE integrate at different genomic sites

The sites at which the GA1 and GA2 ICE insert into the recipient genome have not been previously analyzed. ICE transfer is often a precisely regulated event (Waters and Salyers, 2013), with integration into the recipient genome mediated by an integrase encoded within the ICE. Here, we mapped the termini of GA1 and GA2 ICE and determined their chromosomal integration sites and the sequence specificity of the integrases. For GA1 ICE, integration sites varied widely across genomes. For example, in *B. fragilis* YCH46, the GA1 ICE insertion disrupted a fucosyltransferase gene (BF2787) in a polysaccharide biosynthesis locus (Fig. 5), while in *B. cellulosilyticus* CL06T03C01, the GA1 ICE integrated downstream of an *p-*aminobenzoyl-glutamate transporter gene cluster. Analysis of GA1 ICE flanks indicated that there is little sequence conservation between species at the regions directly upstream and downstream of the ICE. The first gene of the GA1 ICE encodes the integrase of the CTnBST tyrosine recombinase family, which are sequence-selective rather than sequence-specific (Wesslund et al., 2007). Wang *et al*. (Wang et al., 2011) showed that this integrase mediates recombination at sites with the consensus pattern tTnCcAA, where n is any residue, and lower-case letters indicate a single allowed mismatch at one of these two sites. We determined that the GA1 insertion sites in all the genomes we analyzed are flanked by this 7-bp pattern (Fig. 5), explaining our observation that this ICE inserts into diverse locations throughout Bacteroidales genomes. ICE insertion leads to a direct duplication of the 7-bp sequence such that it is found at the upstream and downstream junctions of the ICE. By aligning the flanking genomic regions in *B. fragilis* YCH46 (with a GA1 ICE insertion) and *B. fragilis* 638R (no insertion), we verified that the duplication only spans this 7-bp sequence (Fig 5). This low sequence selectivity may partially explain why GA1 ICEs are widely distributed across *Bacteroides* and *Parabacteroides* species.

**Figure 5.**
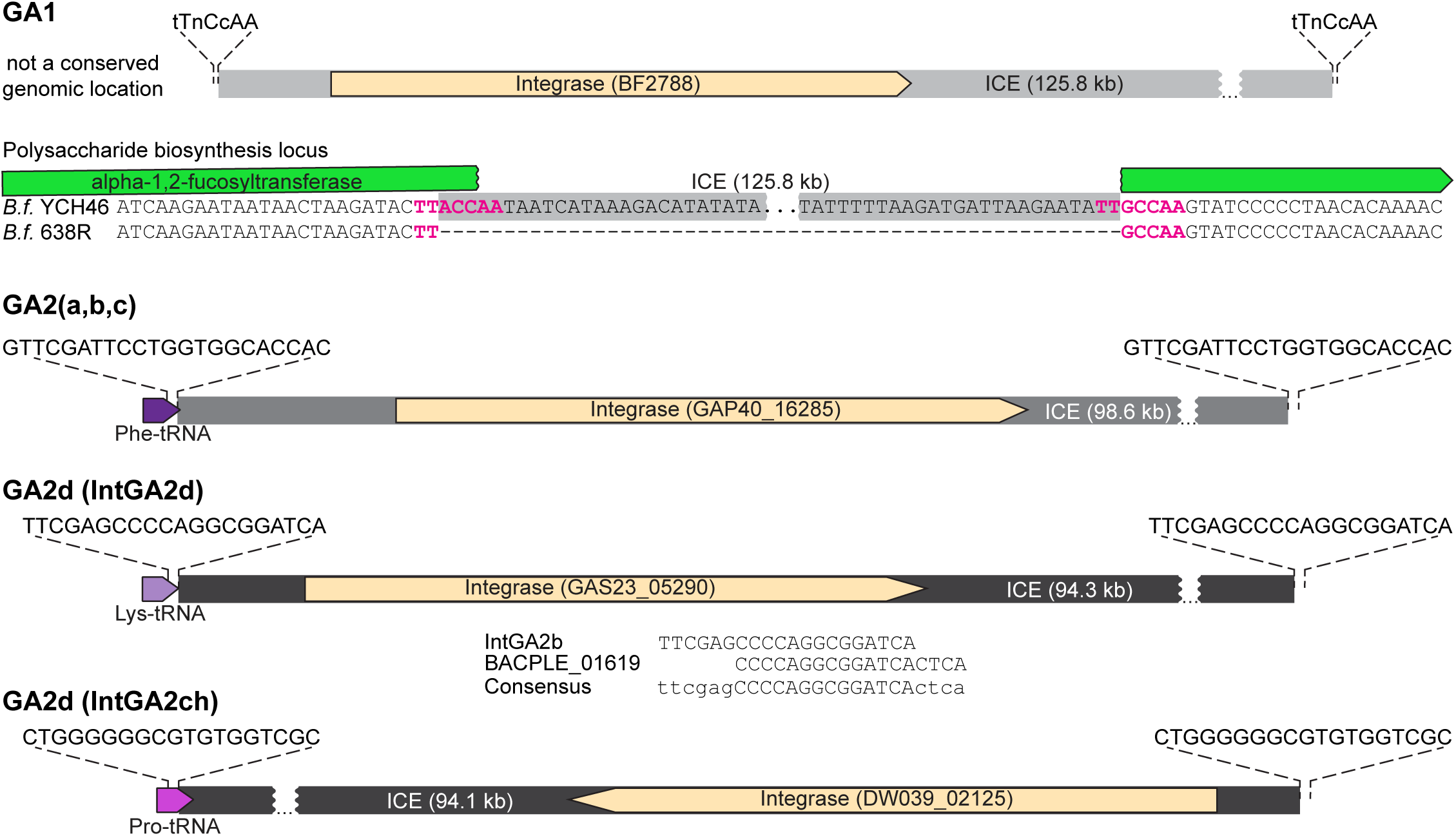
Sequence specificity of the GA1 and GA2 integrases in *Bacteroides* and *Parabacteroides* genomes. **A**. GA1 recombinase and integration site showing the *B. fragilis* YCH46 ICE (BF2788 to BF2921). The insertion of the GA1 ICE in the B. fragilis YCH46 genome relative to the 638R genome is shown. **B**. The recombinase and integration site of the GA2(a,b,c) showing the GA2b region from *B. uniformis* BIOML-A2 as an example (GAP40_16280 to GAP40_16650 and GAP40_07160 to GAP40_07320). This ICE integrate into the Phe-tRNA gene. **C, D**. The recombinases and integration sites of two different GA2d ICE. Panel C shows the GA2d region from *B. uniformis* BIOML-A77 (GAS23_04785 to GAS23_05290) with the IntGA2d integrase. Shown below are the proposed recognition sequence deduced from BIOML-A77 and BIOML-A84 comparison, the sequence reported for *B. plebeius* DSM 17135 (a PUL ice), and the proposed consensus sequence recognized by IntGA2d. This integrase recognizes the sequence shown in Lys-tRNA genes. Panel D shows the GA2d ICE from *B. uniformis* AF39-16AC (DW039_02125 to DW039_02640) which harbors the IntGA2ch recombinase which integrates at a Pro-tRNA gene sequence.

In contrast, GA2 ICE insertions are sequence-specific. For subgroups a, b and c (by far the most abundant), all integrations occurred, with roughly equal likelihood, at either of the two tRNA^Phe^ genes. These ICE are flanked on both sides by the 22-bp sequence GTTCGATTCCTGGTGGCACCAC (Fig. 5). A comparison of the isogenic strains *B. uniformis* BIOML-A2 (GA2 insertion) and BIOML-A3 (no insertion) confirmed duplication of this recognition pattern upon ICE insertion, as it is present only once in the isogenic strain lacking the GA1 ICE. A multiple sequence alignment of the two tRNA^Phe^ regions from species that often carry GA2a/b/c ICE insertions (*B. stercoris, B. uniformis*, and *B. vulgatus*) indicates very little conservation in the 15-bp region downstream of the 22-bp recognition pattern. This downstream region is often AT-rich, but is also AT rich for at least one of the tRNA^Phe^ loci from *B. thetaiotaomicron* and other species that rarely acquire GA2a/b/c, indicating that this high AT region may not be a factor in species prevalence. Moreover, GA2a/b/c insertions do not strictly require an unoccupied tRNA^Phe^ site as we identified one instance (*B. xylanisolvens* CL11T00C41) where a GA2b ICE integrated in tandem downstream of a distinct ICE that integrates at the same site. This GA2b ICE integration follows the 22-bp direct repeat generated by the other ICE’s integration (integrases INE93_03024 and INE93_03154, respectively). The tyrosine recombinase located at the beginning of the GA2a/b/c ICE (IntGA2abc, GAP40_16285) is 81.55% identical at the protein level to the integrase of an α-mannan utilization ICE from *Bacteroides thetaiotaomicron* VPI-5482 (Hehemann et al., 2012). This integrase recognizes the same 22-bp sequence as IntGA2abc, and generates the same direct repeats upon integration. Therefore, the presence or absence of the integrase recognition sequence by itself does not explain the species distribution and abundance of GA2, in particular its relative absence from *B. thetaiotaomicron* and *B. ovatus*. The 22-bp sequence is conserved in *Porphyromonas* species, and while less conserved in *Prevotella* and *Alistipes* species, based on target sequence alone, a significant number of strains that do not contain GA2a/b/c ICE contain the sequence for integration.

The GA2d ICEs do not have a single conserved integrase and therefore do not integrate at a single site. The genomes of seven strains in our collection (Table S1) contain a GA2d ICE. In one isolate, *Bacteroides eggerthii* AF18-1 (from the CGR collection), the GA2d T6SS region is contained on an ICE similar to GA2a/b/c ICE, with the same insertion site described above. For the other strains, the GA2d ICE is unrelated to the GA2a/b/c ICE and has one of two different integrases that dictate their site of integration. The ICE of five strains, *B. vulgatus* BIOML-A119, *B. uniformis* BIOML-A85, *B. dorei* BIOML-A25, *B. plebeius* AM31-10, and *B. vulgatus* VPI-4496.2, inserted into the tRNA^Lys^ locus. By comparing the isogenic strains *B. uniformis* BIOML-A84 (no GA2d ICE insertion) and BIOML-A85 (GA2d insertion) isolated from the same person, we determined that the 20-bp recognition pattern for the integrase of this ICE (IntGA2d, GAS23_05290 in *B. uniformis* BIOML-A77) is TTCG<u>A</u>GCCCCAGGCGGATCA, which duplicates upon ICE insertion (Fig. 5). The ICE also inserted at this site in the other four species, and the sequence repeats on each side of the ICE, with only one mismatch in one flank of the insertion for in *B. vulgatus* VPI-4496.2 (the underlined A in the pattern was substituted by a G). Interestingly, a closely related ICE harboring a porphyran utilization locus (98.57% sequence identity over 48.5% of the core ICE excluding cargo) was characterized in *B. plebeius* DSM 17135. The phage-like recombinase from this ICE is 100% identical to IntGA2d (Hehemann et al., 2012). These authors concluded, based on PCR and sequencing of the products from rare ICE excision events, that the 18-bp recognition sequence for the *B. plebeius* integrase (BACPLE_01619) is CCCCAGGCGGATCACTCA. This pattern does not completely match the direct repeat we identified for IntGA2d (Fig. 5). Some *Bacteroides* integrases are known to tolerate a small number of mismatches in the overlap sequence, which may account for this discrepancy (Wood and Gardner, 2015). Combining these data, it is predicted that this integrase recognizes/duplicates a 24-bp sequence ttcgagCCCCAGGCGGATCActca with some tolerance for mismatches (lower case) outside the core region (capitalized).

Intriguingly, the GA2d ICE from one isolate, *B. uniformis* AF39-16AC, shares lower sequence similarity with the rest of GA2d ICE regions (84.3% average identity) and does not have IntGA2d at the beginning of the ICE. Instead, a 4 kb insertion located at the 3’ end of the ICE harbors a different phage-like recombinase (IntGA2ch), which bears only 24.1% protein sequence identity to IntGA2d and has no characterized orthologs. A similar GA2d ICE with IntGA2ch is present in the chicken isolate *Bacteroides* sp. An19. The 4 kb insertion carrying IntGA2ch is also present in a closely related ICE (96% identical) harboring a different cargo (the *B. dorei* HS1_L_1_B010 ICE up to the EL88_18285 integrase). In all three cases, the ICE insertion occurred at a tRNA^Pro^ locus and is flanked by the 19-bp sequence CTGGGGGGCGTGTGGTCGC.

In sum, these observations highlight that recombination events between related ICEs in the Bacteroidales can lead to changes in genomic target location due to integrase swaps, as well as changes in gene cargo of the ICE. In all examples identified, GA2 ICE integrate in tRNA genes, integrating at Phe-tRNA, Lys-tRNA or Pro-tRNA genes.

Based on these insertion sites, it is not known why GA2 T6SS loci are scarce in species such as *B. thetaiotaomicron, B. ovatus, B. fragilis*, and *B. intestinalis*, while present in more than 30% of the strains of other species such as *B. uniformis, B. vulgatus* and *B. eggerthii*. Whether this bias occurs at the transfer, integration, or maintenance stage is currently unknown. It is possible that these large ICE with their T6SS loci may impose a fitness cost in some species that would outweigh any fitness advantage, precluding their selection and ecosystem sweeps. Alternatively, there may be species-level differences in the ability to properly regulate or selectively silence elements on the ICE or T6SS that may lead to decreased fitness. Successful integration and proper regulation of the complex T6SS machinery into different underlying cell physiologies and environmental niches is expected to be highly variable. Our data suggest that GA1, which are present in numerous species with no obvious bias, may present fewer barriers to their acquisition and maintenance.

### Exclusion of GA2 loci with GA1 and GA3 T6SS loci

Based on our analysis of the distribution of the T6SS genetic architectures and their sweeps within ecosystems, we suspected that the presence of a GA2 T6SS or ICE may preclude the acquisition of a GA1 ICE or vice-versa. In our Bacteroidales isolates genome collection (Table S1), only eight strains were identified as having both a GA1 and GA2 T6SS locus (excluding from this count one of the two nearly-clonal strains *B. vulgatus* AF15-6A and AF16-11). In addition, *B. fragilis* strains frequently harbored both a GA3 and GA1 T6SS locus (23 strains), but rarely both a GA3 and GA2 T6SS loci (1 strain). Interestingly, two of the eight strains with both a GA1 and GA2 ICE have disruptions in the T6SS loci. In *B. vulgatus* BIOML-A11 (and all its other 79 isogenic co-isolates) the *tssC* gene from GA1 is frameshifted by 2 bp at codon 148 of 460 (Fig. 6). In strain *B. uniformis* BIOML-A5, isolated from the same subject (“am”), the GA1 is intact but the *vgrG* gene has a 5 bp frameshift at codon 346 of 603 (Fig. 6).

**Figure 6.**
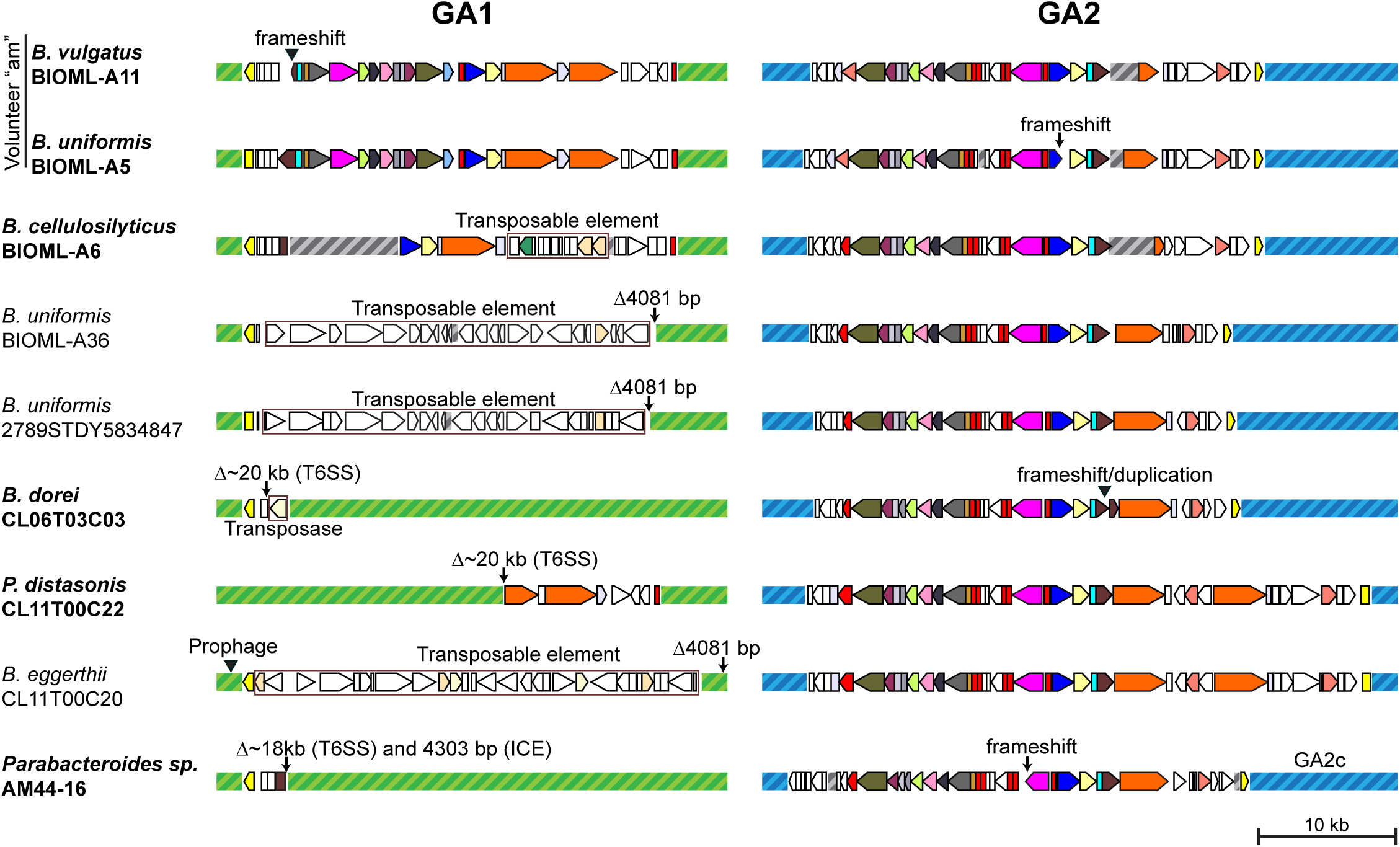
Schematic of genomes that have a GA1 and a GA2, where one of the regions is disrupted. Corresponding genomic coordinates for each region shown are included in Table S6. GA1 backbones are shown as hatched green boxes and GA2 backbones are indicated by hatched blue boxes. All GA2 regions shown are of the GA2b subtype except where indicated. Grey hatched boxes indicate gaps in contigs or contigs that were joined manually. Color legend as specified in Figure 1. Strains with bolded names have disrupted T6SS regions, whereas non-bolded correspond to the same ICE carrying a different cargo. Insertions are indicated by filled triangles, deletions by arrows. Brown boxes indicate the bounds of a transposable element insertion or recombination event that replaced the cargo.

ICE entry exclusion is a relatively common phenomenon in Gram-positive bacteria and proteobacteria, where a gene encoded within the ICE acts to prevent the acquisition by the cell of the same or similar ICE (Avello et al., 2019). However, this is an unlikely explanation for GA1/GA2 exclusion as there is little sequence conservation between them. To determine whether the main determinant of GA2 exclusion is the ICE or the cargo, we conducted a similarity search for GA1 and GA2a-d ICEs (rather than the T6SS locus) in all Bacteroidales isolate genomes listed in Table S1 and in the community isolate datasets (Table S3). We identified five additional instances of GA1 and GA2 ICEs present in the same strain where one of the T6SS was frameshifted, truncated or replaced by a transposable element (Fig. 6, Table S6). These observations suggest a functional and active T6SS may directly prevent acquisition of a second T6SS (akin to an entry exclusion mechanism, for instance by killing the donor cell via T6SS toxins), such that it only happens if the first T6SS is already disrupted. Alternatively, there may be a fitness disadvantage associated with having an intact GA2 T6SS together with a GA1 or a GA3, such that when they co-occur there could be a selective pressure for disruption of one of the loci. However, we were unable to identify any salient features in the T6SS regions of the six strains with both intact GA1 and intact GA2 T6SS loci except that five are from the CGR dataset from Chinese individuals. Notably, for these strains, there was no protein sequence overlap in toxin/immunity genes between the two GA regions nor did they contain acquired interbacterial defense islands carrying the cognate immunity genes to the toxins in their GA1/GA2 (Ross et al., 2019). It is possible that one or both of the T6SS may be silenced, in particular in the case of strain AF15-6A (and closely related AF16-11) which carries a GA1, a GA2b, and a GA2c.

Additionally, we found three instances of a GA2 ICE in a strain carrying an ICE closely related to that of GA1 (99.3% identity and 95.2% coverage) but with a different cargo instead of the T6SS (Fig. 6, Table S6). A comparison of the ICE between strains with only one ICE versus the 15 strains where the two co-occur (including the three with different cargo) showed nothing remarkable about the GA2 locus in strains with GA1. In contrast, this comparison allowed us to identify a gene of interest in the GA1 ICE that may be involved in exclusion. This gene (BF2858 in *B. fragilis* YCH46) is absent or disrupted in 11 of the 16 strains (73.3%), due to independent instances of transposase insertions. In contrast, the gene is disrupted in 31.2% of the 125 non-redundant strains with only a GA1 ICE in tables S1 and S3 (p = 0.0049 on Fisher’s exact test, comparing disrupted vs. non-disrupted and GA1 alone vs. GA1 and GA2). The gene, which we named MeGA1, encodes for a 1660-residue protein containing both an *N*^6^-adenine methylase domain and an SNF2 helicase domain. This architecture is reminiscent of type I/III restriction modification systems, although for MeGA1 the helicase domain does not have sequence signatures predictive of endonuclease activity (Schwarz et al., 2013). Recently, another type of phage-defense system (DISARM) was described which also carries an adenine DNA methylase gene and a SNF2 helicase gene. These proteins, together with another helicase and a DUF1998 domain protein, provide protection from lytic and lysogenic phage infection in *Bacillus subtilis* through an unknown mechanism (Ofir et al., 2018). The association of the defect in the MeGA1 gene with strains that have both a GA1 and GA2 ICE is intriguing, but it is unknown how such a product would participate in the exclusion of an element acquired in single-stranded form. Nevertheless, the observed GA2 ICE exclusion may therefore be the result of two distinct factors: ICE exclusion of GA2 by an unknown property of MeGA1 and T6SS exclusion. The latter could be caused by the difficulty of a single cell simultaneously maintaining two complex machines that may cross-talk and are each expensive to replicate and fire. Such a combination may therefore only rarely reach fixation in a population.

### Metagenomic analyses of T6SS

Analyses of genome sequences has been informative in many regards, but do not reveal the prevalence of these T6SS loci in the metagenomes of human populations. Among Bacteroidales species, human gut microbiomes tend to be dominated largely by *Prevotella* or by *Bacteroides/Parabacteroides* (Arumugam et al., 2011; Costea et al., 2018). We analyzed 15 different human gut metagenomic datasets including metagenomes from 1767 individuals to identify the global distribution of the three different T6SS GAs and the five different GA2 subtypes. In these composite metagenomes, GA2 T6SS loci are the most abundant of the three T6SS GA, where their abundance in *Bacteroides*/*Parabacteroides* dominated communities such as the Japanese and US datasets is 67% and 43% of metagenomes, respectively. GA1 T6SSs are also very prevalent in *Bacteroides/Parabacteroides* dominated communities, present in 50% of Japanese metagenomes and 45% of US gut metagenomes. Other populations such as Mongolians and Fijians have a greater number of metagenomes with GA2 T6SS compared to GA1 T6SS loci (Table S7, Fig 7). Among the different subtypes of GA2, there are associations with different populations. For example, 74% of the GA2 T6SS in the Mongolian dataset are subtype GA2a, with only two gut metagenomes containing a GA2b locus. In contrast, the GA2b subtype dominates in the population comprising the Japanese dataset (65% of GA2) and GA2c dominate in the Fiji dataset (65% of GA2 loci). The GA2d subtype are only present in 25 metagenomes with a global rather than clustered distribution. No GA2e were detected in any of the 1767 human gut metagenomes queried, further supporting that this GA2 subtype is largely present in Bacteroidales strains of animals.

**Figure 7.**
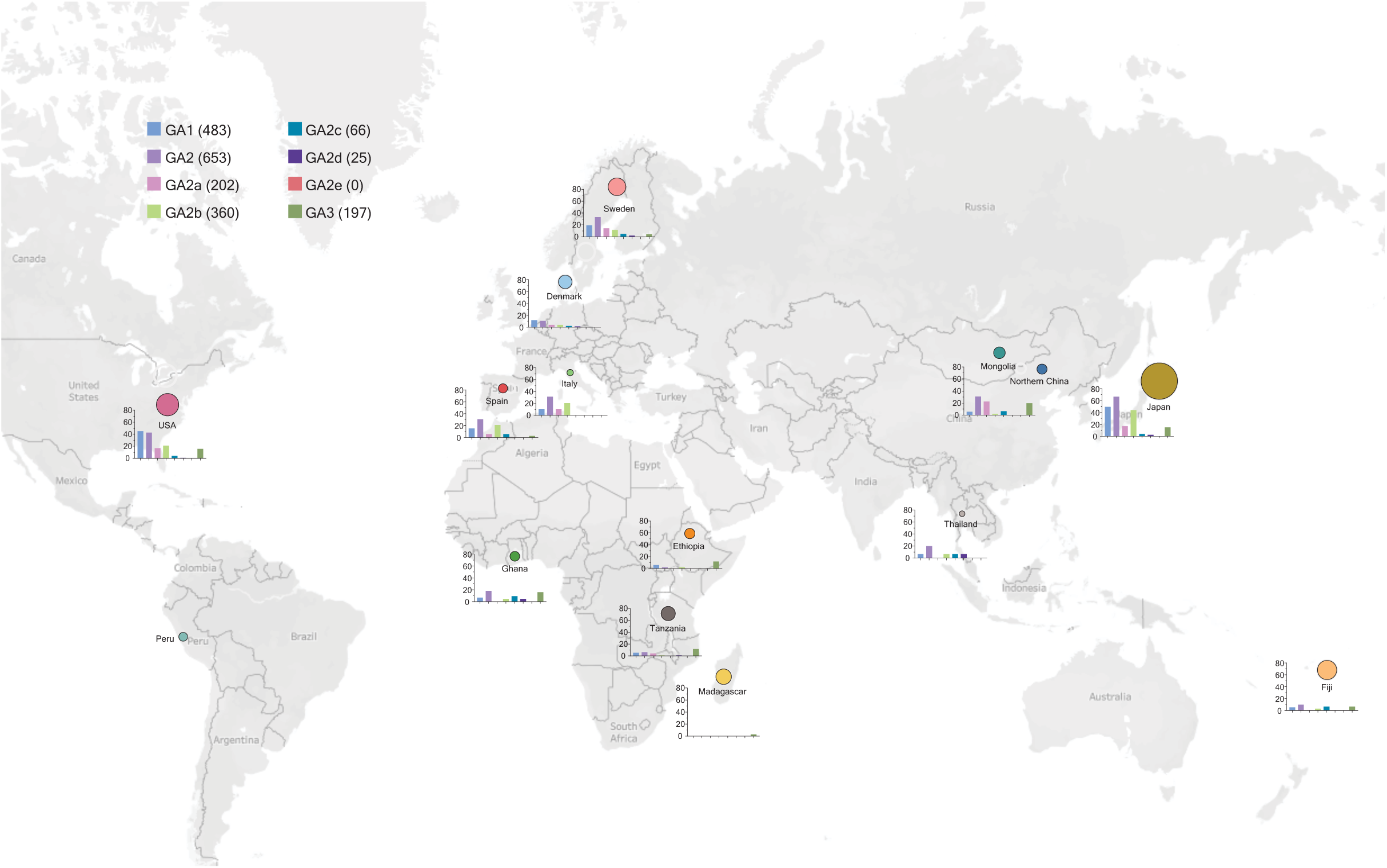
Distribution the T6SS GA loci and subtypes in human gut metagenomes from different human populations. The percentage of metagenomes with each GA and GA2 subtype of the total metagenomes for each population is shown by bar graph. The size of each population sampled is designated by the size of the circle. No GA loci were detected in the Peru metagenomic dataset. The exact numbers for each metagenomic dataset are reported in Table S7. World map from © 2020 Mapbox © OpenStreetMap

### Other MGEs sweep through communities but most to a lesser extent

To identify other mobile genetic elements (MGE) in gut Bacteroidales and if they undergo multi-species sweeps within a person’s microbiota as commonly as those of GA1 and GA2 ICE, we scanned all four community isolate datasets (CL, BIOML, UK and CGR collections) for MGEs (4 kb or larger) that are identical in at least three different species within a community. We identified multi-species sweeps of at least one MGE in 67 out of 91 (73.6%) human gut communities. We identified 389 occurrences of MGE sweeps, which clustered into 177 MGE groups based on a threshold requiring 80% identity with at least 80% of the element shared within a group (Table 2, Table S8, Table S9). Many of these MGEs are present in communities where they did not sweep through multiple species, and therefore were not counted. The majority of the MGEs only swept through one (96) or two communities (59), while 14 swept in three different communities.

Only nine MGEs, including GA1 and GA2b/c ICEs, were found to sweep through multiple species in four or more communities with a fully conserved architecture (Table S8, Table S9). From these conserved and commonly sweeping MGEs, one ICE, CTn341 (Bacic et al., 2005) carrying a tetracycline resistance gene, was ubiquitous with several variants. At least one of its five variants are present in 94% of communities and swept in multiple species in 44%. Other MGEs are less ubiquitous but nonetheless show widespread sweeps, albeit at a lesser frequency than GA1. There are numerous predicted phenotypes encoded on these MGEs. One contains genes encoding putative vitamin B12, hemin, and iron transporters. Three others carry genes encoding putative antibiotic resistance determinants. A 98 kb conjugative megaplasmid, which we named pMMCAT, is transferred to many species in six communities and carries an extracellular or capsular polysaccharide biosynthesis gene cluster and a locus with fimbriae genes. Interestingly, plasmids consisting of smaller regions of this megaplasmid (one without the EPS locus and one without the conjugation machinery and the EPS locus) are also observed sweeping in three communities each in the CGR collection. These data highlight the abundant and rapid evolution of human gut Bacteroidales species via mobile genetic elements encoding numerous functions likely related to microbe-microbe and microbe-host interactions.

Rearrangements in these MGE were common, such that different MGEs are partially overlapping, indicating occasional loss, gain or swapping of genes. Other studies of inter-species horizontal gene transfer showed a similar trend in that clusters of functional genes (cargo) and their encompassing mobilization vehicles often undergo recombination and other rearrangements, such that MGE architectures are not conserved (Brito et al., 2016; Jiang et al., 2019). For instance, one cluster of genes for the breakdown of a glycogen-like molecule was found to be carried by three different MGEs (clusters 12, 13 and 91 in Table S8).

We analyzed the metagenomic dataset for the presence of the frequently sweeping MGEs clusters (clusters 1,4,6,7,8,10, 12) (Table S10). Cluster 1 (CTn341 and related ICEs), was strikingly ubiquitous across all sampled populations, locales, and lifestyles, including individuals with very low *Bacteroides* and *Parabacteroides*. Individuals where this ICE is absent are very rare. This same pattern is followed, to a lesser extent, by the cluster 6 Integrative Mobilizable Element (IME), which carries a putative extended spectrum β-lactamase. This type of MGE lacks conjugation machinery and relies on other MGEs for transfer. Interestingly, both of these MGEs, which carry antibiotic resistance genes, are present in many Bacteroidales genera and are ubiquitous in recent isolates. In contrast, the other MGEs are less ubiquitous in genomes and metagenomes, with differences in abundance related to geography and lifestyle (Table S10). pMMCAT, which only transfers among *Bacteroides* and *Parabacteroides*, is ubiquitous (>70% abundance) in populations from USA, Japan and Thailand. It is also present at substantial frequencies (22-70%) in all the other studied populations except for Madagascar, hunter-gatherers in Tanzania and Peru, and an agricultural community in Peru. Clusters 7 and 12 follow roughly similar distribution patterns, largely correlating with *Bacteroides* and *Parabacteroides* abundance. These widespread and frequently sweeping MGEs, akin to GA1 and GA2, seem to have favored architectures, with infrequent rearrangements. In contrast, most other MGEs in the Bacteroidales are subject to frequent rearrangements. One caveat to note is that our MGE sweep analysis was conducted using four datasets from industrialized lifestyles, largely from Bacteroides-dominated communities. Analyses of HGT events in *Prevotella*-dominated non-industrialized populations indicated less frequent HGT (Groussin, 2020) and may show a different repertoire of MGEs and different frequency of multi-species sweeps.

Through the genomic analyses of more than 1500 Bacteroidales strains, many co-resident in the same individual, combined with analyses of nearly 1800 gut metagenomic data sets of diverse global populations, we have uncovered numerous properties of the T6SS of the gut Bacteroidales. Previously thought to be rarely present in the human gut microbiota, we show that GA2 T6SS loci are the most prevalent of the three GA in the human gut metagenomes analyzed here and that T6SS of GA2 segregate into five subtypes, one of which is largely restricted to animals. GA2 subtypes in many cases demonstrate population clustering with GA2b the most abundant in Japan, but rarely present in the Mongolian population sampled where the GA2a subtype dominates. GA1 ICE acquisition and spread may be further facilitated by the low sequence selectivity necessary for integration. Most importantly, this study reveals the extensive intra-ecosystem transfer of these ICE to co-resident members of the gut microbiota, suggesting a potential community benefit by multi-species acquisition. This type of globally conserved architecture is rare for MGEs that sweep in multiple species within an individual. This observation is especially interesting for GA1 and GA2, given that the T6SS variable regions harboring the effectors are frequently recombined, yet the general architecture is preserved.

## Materials and Methods

### Creation and curation of a Bacteroidales isolate genome collection

All genomes classified by NCBI in April 2020, as belonging to the order Bacteroidales, excluding genomes that were suppressed or considered anomalous, and excluding genomes flagged as derived from metagenomics studies, surveillance projects, or environmental sources, were identified by an Entrez query. This initial set of 2,324 genomes was further curated to reduce clearly redundant genomic sequences (e.g. the same strain sequenced multiple times, including equivalent entries from strain repositories), reduce longitudinal samples comprising the same strain isolated from the same individual multiple times, and to remove duplicate entries from both the GenBank and Refseq repositories, retaining the RefSeq sequence. Finally, genomic sequences that were assembled from metagenomics studies despite not being flagged as such by NCBI and those genomes not identified to at least the genus level were removed. The final set comprises 1,434 Bacteroidales genomes (Table S1).

### PCR screening for GA2 T6SS

We screened 136 *Bacteroides* and *Parabacteroides* isolates collected from four volunteers in the Comstock lab strain collection previously isolated under a protocol approved by the Partners Human Research Committee IRB and complied with all relevant federal guidelines and institutional policies (Zitomersky et al., 2011a). Each strain was spotted on a BHIS plate and grown anaerobically. A very small amount of cell material was collected from the colony, resuspended in 50 µl water, boiled for 10 minutes and centrifuged. 1 µl from the supernatant was used for a 20 µl PCR reaction with Phusion polymerase (NEB), using primers oLGB21 (TGGGAGCAAGTTTTCTGAATTTGG) and oLGB22 (TGTTCTCCTGCGCTACATAATCGTATC) for the conserved region, and primers oLGB27 (CKTGAATTGAACATCCATTCCAR, where K = G,T and R = A,G) and oLGB28 (GATCCAGTGGATGCTGGATG) for the variable region (Figure 3A). The annealing temperatures were 54°C for the conserved region and 64.5°C for the variable region, with extension times of 50s and 1m45s, respectively. For the variable regions, PCR bands were purified from an agarose gel and sequenced using Sanger sequencing.

### DNA extraction, sequencing and genome assembly

The genomic sequencing of bacterial strains isolated from human fecal samples was approved by the Partners Human Research Committee IRB and complied with all relevant federal guidelines and institutional policies. Strains (Table S2) were grown anaerobically in basal medium as described previously (Pantosti et al., 1991) to an OD600 of ∼0.8. DNA was recovered using a CTAB/NaCl DNA extraction protocol followed by sodium acetate/ethanol precipitation. SMRT sequencing was carried out at the University of Maryland’s Institute for Genome Sciences Genomics Resource Center, who performed the initial quality control, library preparation, and sequencing of the genomes using PacBio Sequel v3 SMRTcell technology. The genomes were assembled separately using Falcon/Unzip 1.2.0 (Chin et al., 2016) and Flye 2.8.2 (Kolmogorov et al., 2019), then reconciled using Flye. In cases where unresolved bubbles where still present (*B. uniformis* CL06T03C05, *B. dorei* CL06T03C03, *B. cellulosilyticus* CL06T03C01, *B. xylanisolvens* CL11T00C03, *B. eggerthi*i CL11T00C20) as assessed by Bandage (Wick et al., 2015), the two assemblies were also reconciled with a Canu 1.8 (Koren et al., 2017) assembly. Remaining ambiguities due to invertible DNA regions were fixed manually, using the inverted repeat regions as bounds and available complete reference genomes as guidance. Reconciled assemblies were polished using GCpp 2.0.0 (Pacific Biosciences). Genomes from *B. ovatus* CL03T12C18, *B. vulgatus* CL11T00C01 and *B. xylanisolvens* CL03T12C04, where multiple contigs were obtained or bubbles remained unresolved, were scaffolded using Ragout 2.3 using complete genomes from the same species as reference (Kolmogorov et al., 2018). Genes were called using Prodigal 2.6.3 (Hyatt et al., 2010) and annotation was performed using a customized version of Prokka 1.14.6 (Seemann, 2014). Small contigs were classified as plasmids based on circularization during assembly and using PlasFlow (Krawczyk et al., 2018). The genomes were submitted to GenBank and assigned BioProject accession number PRJNA669351.

### Community collections

In addition to our own longitudinal isolates (see below), we utilized three other publicly available isolate sequence sets in our analysis of intra-community DNA transfer. Genomes from these sets were included if they were identified as belonging to the order Bacteroidales and comprised at least three different species isolated from the same individual. These included: 967 genomes from BioProject accession PRJNA544527 (BIOML collection, (Poyet et al., 2019) representing 23 species of 4 genera collected from 10 individuals; 380 genomes from BioProject accession PRJNA482748 (CGR collection, (Zou et al., 2019) comprising 30 species of 6 genera collected from 71 individuals; 27 genomes from BioProject accession PRJEB10915 (UK collection, (Browne et al., 2016) representing 13 species from 4 genera sampled from 5 individuals; and finally the CL collection comprised 42 genomes from 18 species of two genera collected from five volunteers (Table S3). It included the 23 genomes sequenced in this study (Table S2, BioProject accession PRJNA669351) plus GenBank accessions GCA_001640865.1, GCA_000307345.1, GCA_000273015.1, GCA_000273035.1, GCA_000273055.1, GCA_000273175.1, GCA_000273235.1, GCA_000273235.1, GCA_000307395.1, GCA_000307375.1, GCA_001535615.1, GCA_001535595.1, GCA_000269545.1, GCA_001535605.1, GCA_000273295.1, GCA_000307435.1 and GCA_000307495.1. These 1,530 genomes were used to create blast databases for intra-community DNA transfer analysis.

### Distribution of T6SS genetic architectures in genome sequences of Bacteroidales isolates

The T6SS loci found in species of the Bacteroidales order fall into three genetic architectures (GA). All three of these GAs contain structural genes (i.e. the genes named with the *tss* nomenclature) which are consistent as to order and identity within a GA. They also each contain variable regions encoding such things as toxin and immunity proteins, RHS proteins, and evolved TssD proteins. These variable regions would complicate analysis of the distribution of these GAs in both metagenomic datasets and in sequenced isolates. We therefore made DNA-level concatemers by removing the sequence encompassing the variable region(s) from a representative of each GA, and used these as templates for metagenomic read mapping (see above) and as queries for analysis of sequenced isolates.

The curated set of 1,452 Bacteroidales genomes (see above and Table S1) were processed into nucleotide and protein blast databases using Blast v. 2.10.0+, and the T6SS concatemer templates were used as queries using blastn. The results were parsed and the presence or absence of T6SS GA(s) was determined for each genome (see Table S1). The majority of the results for all seven queries (GA1, five GA2 subtypes, and GA3) were unambiguously clear (e.g. a sequence span detected in an isolate that was greater than 95% identical covering 95% or more of the query). More ambiguous returns with less coverage or lower DNA identity were inspected manually by retrieving the segment(s) involved, usually including some flanking DNA, from the isolate’s genome sequence, and scaffolding hits spanning multiple contigs from heavily fragmented genome sequences. Further comparative analyses were also performed between all concatemer templates and all subject sequences using Clustal Omega v 1.2.4 ((Sievers et al., 2011)) under Linux (CentOS v 8).

### Phylogenetic trees

Phylogenetic analyses were based on 16S sequences acquired from public repositories such as RDP and Silva. Representative 16S sequences of Bacteroidia identified as type strains by JGI were analyzed for phylogeny using MEGA X (Kumar et al., 2018). After alignment with Clustal as implemented by MEGA X, the Maximum Likelihood trees were created under the General Time Reversible model (GTR), using a discrete gamma distribution to model evolutionary rate and a rate variation model that allowed some sites to be evolutionarily invariable. Initial trees were obtained via Maximum Parsimony method. The trees shown are the bootstrap consensus trees inferred from 500 replicates. To build the tree of GA2 T6SS regions, sequences were aligned in Clustal Omega and the tree was computed using RAxML (Stamatakis, 2014)using the GTR model, ML estimate of stationary base frequencies, gamma distribution to model among-site rate heterogeneity and a bootstrapping cutoff of 0.03.

### Analysis of T6SS transfer events within co-resident strains

T6SS regions in the community genome collection were identified as specified previously. To determine if T6SS regions were identical within isolates from the same community, regions were retrieved with 10 kb flanking regions on both sides and aligned using Clustal Omega. Regions were determined to be subject to a recent transfer event if they were 99.95% identical over the complete span of the T6SS, including the variable regions.

### Identification of other mobile genetic elements that transfer within individual microbiomes

A separate nucleotide blast database was created for each community set. Each genome from the community was compared against the database using blast, with cutoffs of 99.95% identity and alignment length of 4000 bp. Hits were retained if they were present in three or more isolates of different species. Redundancy between hits was reduced using cd-hit (Fu et al., 2012). Since many of the genomes analyzed are heavily fragmented (and to by-pass the abundant transposase insertions in Bacteroidales genomes), non-redundant hits were blasted against each other and joined using Flye if there was more than 4000 bp overlap at 99.95% identity. The resulting regions, together with those that didn’t require joining, were once again blasted against the community genome database, to verify that each was still present in at least three different species. Genomic regions that fulfilled these conditions were considered to be MGEs subject to recent (i.e. during the lifetime of the subject) within-host multi-species sweeps. MGEs were clustered into similar groups using blast, with an 80% identity cutoff and requiring that the alignment covers 80% of the larger fragment.

### Metagenomic datasets, read mapping, and compositional profiling

Fifteen publicly available metagenomic datasets were utilized in our investigations into the prevalence and distribution of various sequences of interest. Briefly, these sets collectively comprised sequencing reads from 1,767 individuals from geographically diverse regions of the world, and encompassed varying ethnic, cultural, age, gender, health, and lifestyle groups. The metagenomics read sets (see Table S7) were downloaded from the European Nucleotide Archive using Aspera. Tools from the BBMap ver. 38.86 (http://sourceforge.net/projects/bbmap) collection of analysis utilities were used to map reads from these sets to sequences of interest. Though the five GA2x T6SS regions are demonstrably different from one another, they do share a somewhat high level of sequence similarity that might influence short read mapping results. Thus, for these analyses, BBsplit was used, as it maps reads to multiple references simultaneously and, in the case of ambiguity (the read maps to more than one template), will determine the best match and count that read only once. Other mapping analyses were ambiguous matches were not an issue utilized BBmap.

## Acknowledgements

This work was funded by Public Health Service grants R01AI120633 and R01AI093771 from the NIH/National Institute of Allergy and Infectious Diseases to LEC. PacBio genome sequencing was discounted through the Genomics Resource Center at the University of Maryland Microbial Genomics SMRT Grant competition to LGB. The funders had no role in study design, data collection and analysis, decision to publish, or preparation of the manuscript.

## Author contributions

LGB – PCR and genome sequencing and assembly of CL strains/genomes, ICE integration analyses, transfer and sweep analyses, MGE identification and sweep analyses.

MJC - Assembly and curation of strain collections, identification of five GA2 subtypes, strain collection and metagenomic dataset analysis. LGB, MJC and LEC conceptualized the study and wrote the paper.

---

**Table S1**. Prevalence of T6SS regions in a non-redundant collection of sequenced Bacteroidales genomes.

**Table S2**. Prevalence of T6SS regions in the different Bacteroidales species.

**Table S3**. Prevalence of GA1 and GA2 T6SS in the CL, BIOML, CGR and UK strain collections.

**Table S4**. Genomic coordinates for the GA1 T6SS regions in the CL, BIOML, CGR and UK culture collections.

**Table S5. A**. Complete GA1 and GA2 sweeps in cases where only one strain per species is present in the community

**Table S6. A**. Strains with two GA ICE integrations including at least one GA2. **B**. Genomic coordinates of GA ICE in strains with two integrations including at least one GA2.

**Table S7**. Presence of T6SS loci in metagenomic datasets.

**Table S8**. Common mobile genetic elements (MGEs) that carry out multi-species sweeps in four or more Bacteroidales communities.

**Table S9**. Mobile genetic elements that carry out multi-species sweeps in Bacteroidales communities.

**Table S10**. Metagenomic distributions of MGEs that commonly sweep in multi-species of a community.

**Figure S1.**
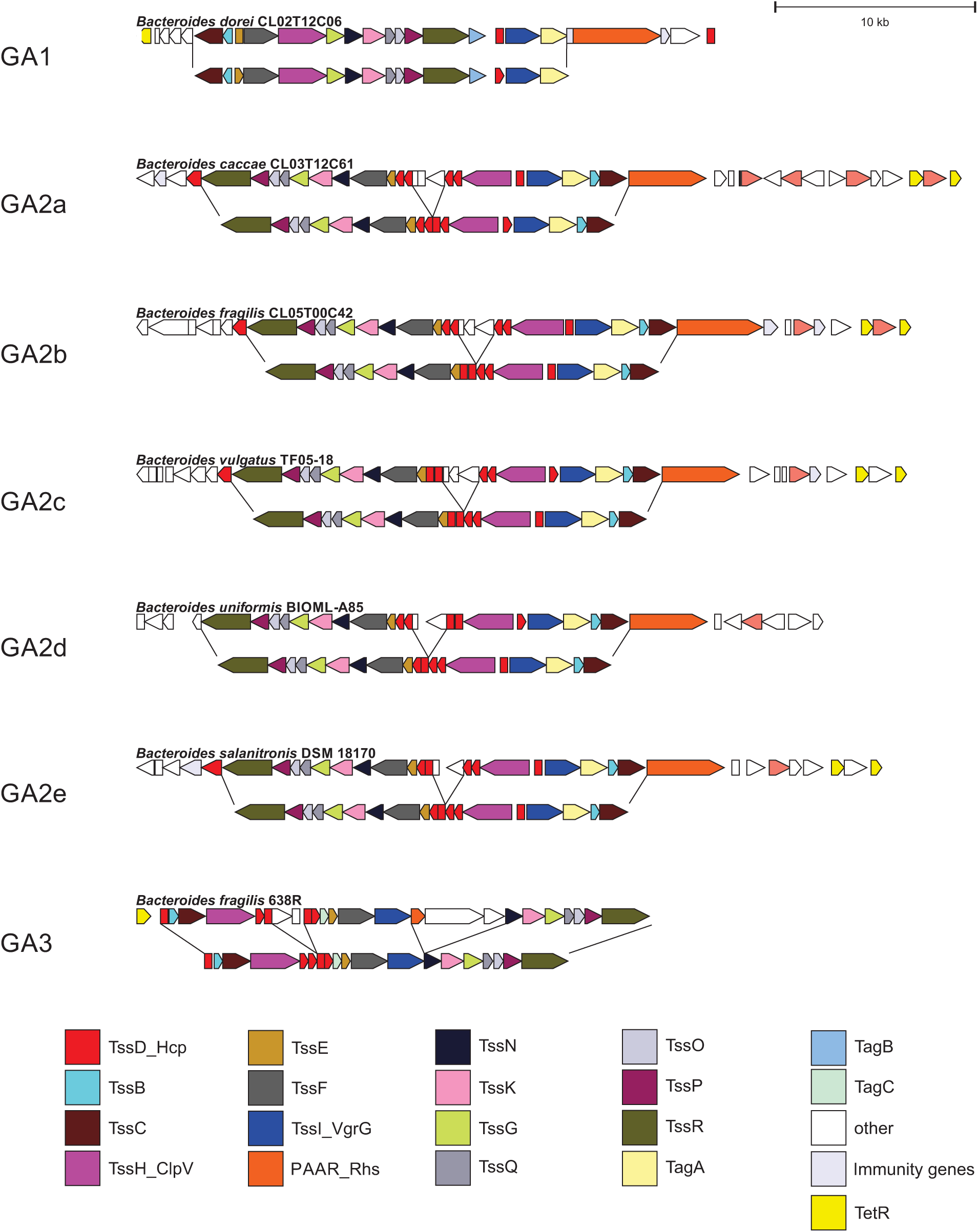
Concatamers of the various gut Bacteroidales T6SS genetic architectures. The bottom gene map of each pair shows the Concatamers that were created after removing all genes that diverge within the same genetic architecture. These Concatamers were used to query the various datasets analyzed in this study.

**Table S5A.** Complete GA1 and GA2 sweeps in cases where only one strain per species is present in the community (For this table we only included strains with at least 4 related isolates).

| Volunteer | Taxonomy | # related isolates | GA1 | GA2 |
| --- | --- | --- | --- | --- |
| af | *Bacteroides caccae* | 4 | y |  |
| af | *Bacteroides fragilis* | 22 | y |  |
| av | *Bacteroides fragilis* | 5 | y |  |
| bf | *Bacteroides fragilis* | 9 | y |  |
| cx | *Bacteroides ovatus* | 22 | y |  |
| cx | *Bacteroides uniformis* | 6 | y |  |
| af | *Bacteroides stercoris* | 4 |  | a |
| am | *Bacteroides fragilis* | 68 |  | b |
| am | *Bacteroides xylanisolvens* | 10 |  | b |
| am | *Parabacteroides merdae* | 11 |  | b |

**Table S5B.**
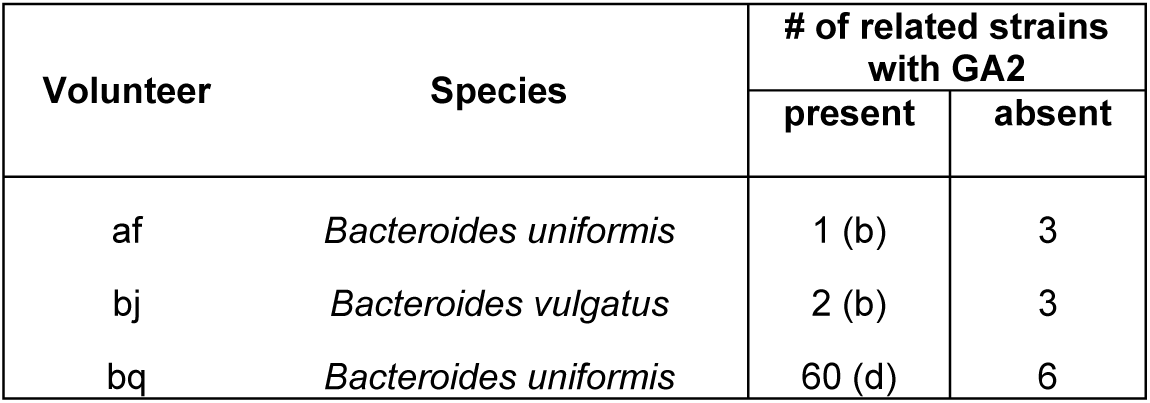
Partial GA2 sweeps in cases where only one strain per species is present in the community.

**Table S5C.** GA1 and GA2 sweeps in cases where more than one unrelated strain per species is present in the community. Isolates were classified into strain groups, each designated by a different letter.

| Volunteer | Strain | Taxonomy | Strain group | No. isolates in group | GA1 | GA2 |
| --- | --- | --- | --- | --- | --- | --- |
| af | BIOML-A15 | *Bacteroides ovatus* | A | 5 |  |  |
| af | BIOML-A13 | *Bacteroides ovatus* | B | 30 |  | c |
| af | BIOML-A1 | *Parabacteroides merdae* | A | 8 | y |  |
| af | BIOML-A3 | *Parabacteroides merdae* | B | 3 |  | b |
| af | BIOML-A4 | *Parabacteroides merdae* | C | 1 | y |  |
| am | BIOML-A38 | *Bacteroides ovatus* | A | 89 |  | b |
| am | BIOML-A84 | *Bacteroides ovatus* | B | 1 |  | b |
| am | BIOML-A110 | *Bacteroides ovatus* | C | 2 |  | truncated |
| am | BIOML-A46 | *Bacteroides vulgatus* | A | 6 |  | b |
| am | BIOML-A11 | *Bacteroides vulgatus* | B | 76 | y, 2 truncated | b, 4 truncated |
| av | BIOML-A11 | *Bacteroides uniformis* | A | 2 | y |  |
| av | BIOML-A12 | *Bacteroides uniformis* | B | 5 |  |  |
| av | BIOML-A106 | *Bacteroides vulgatus* | A | 2 | y |  |
| **av** | **BIOML-A97** | ***Bacteroides vulgatus*** | **B** | **7** | **y** |  |
| **av** | **BIOML-A105** | ***Bacteroides vulgatus*** | **B** | **1** |  |  |
| bf | BIOML-A153 | *Bacteroides ovatus* | A | 1 | y |  |
| bf | BIOML-A154 | *Bacteroides ovatus* | B | 5 |  |  |
| bf | BIOML-A19 | *Bacteroides uniformis* | A | 1 | y |  |
| bf | BIOML-A20 | *Bacteroides uniformis* | B | 4 |  |  |
| bf | BIOML-A24 | *Bacteroides uniformis* | C | 10 |  | a |
| bf | BIOML-A26 | *Bacteroides uniformis* | C | 1 |  | a, truncated |
| bq | BIOML-A26 | *Bacteroides dorei* | A | 18 |  | b |
| bq | BIOML-A17 | *Bacteroides dorei* | B | 1 |  |  |
| bq | BIOML-A25 | *Bacteroides dorei* | C | 1 |  | b, d |
| cx | BIOML-A16 | *Bacteroides stercoris* | A | 6 |  |  |
| cx | BIOML-A18 | *Bacteroides stercoris* | B | 1 | y |  |

**Table S8.**
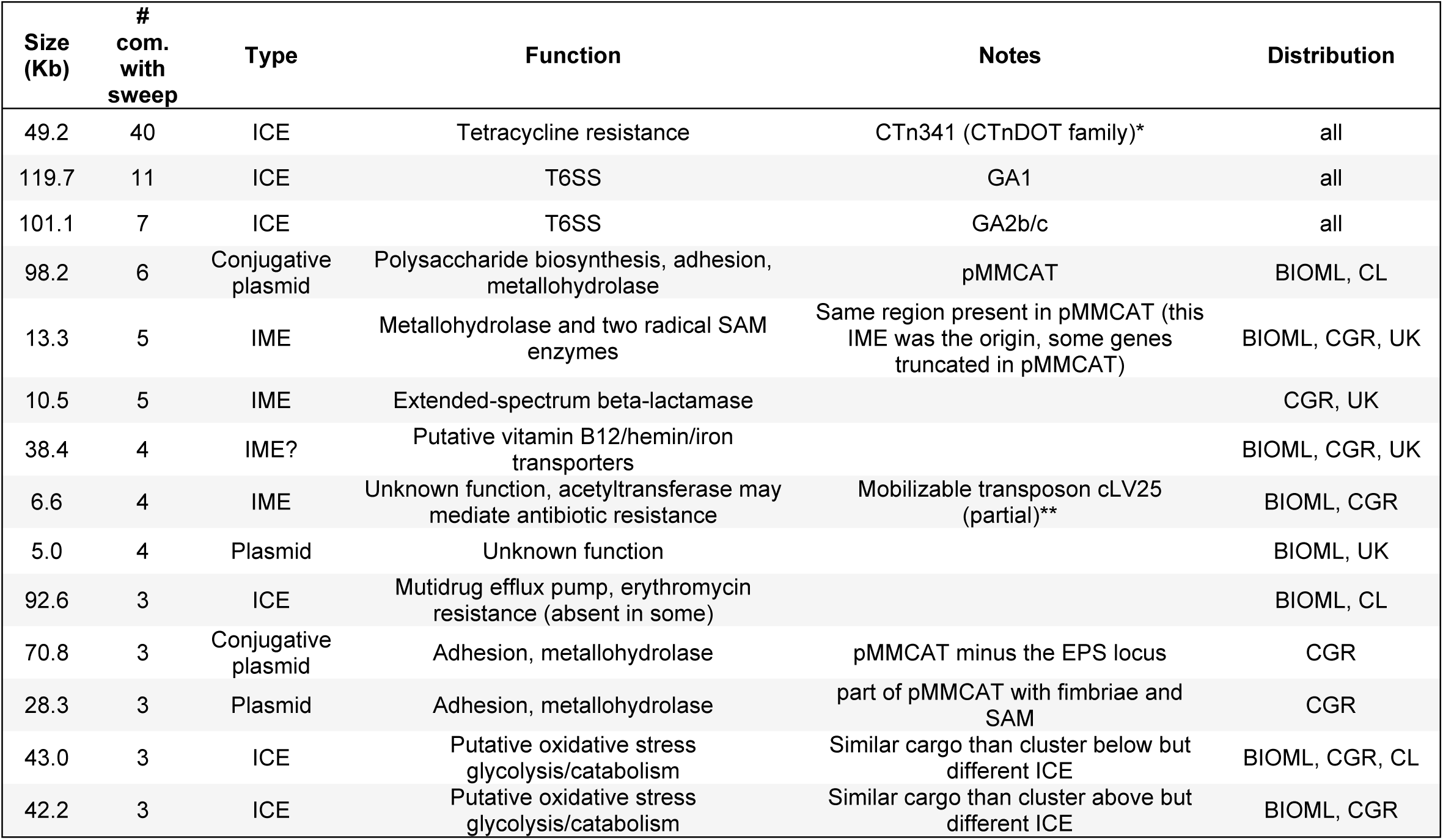
Common mobile genetic elements (MGEs) that carry out multi-species sweeps in four or more Bacteroidales communities. Some additional MGEs that sweep in three communities are shown due to their relevance discussed in the text. All 177 types of MGEs detected to carry out multi-species sweeps are listed in Table S9. *{Bacic, 2005 #2166}, **(Bass and Hecht, 2002)

## References

Arumugam, M., Raes, J., Pelletier, E., Le Paslier, D., Yamada, T., Mende, D.R., Fernandes, G.R., Tap, J., Bruls, T., Batto, J.M., et al. (2011). Enterotypes of the human gut microbiome. Nature 473, 174–180.

Avello, M., Davis, K.P., and Grossman, A.D. (2019). Identification, characterization and benefits of an exclusion system in an integrative and conjugative element of Bacillus subtilis. Mol Microbiol 112, 1066–1082.

Bacic, M., Parker, A.C., Stagg, J., Whitley, H.P., Wells, W.G., Jacob, L.A., and Smith, C.J. (2005). Genetic and structural analysis of the Bacteroides conjugative transposon CTn341. J Bacteriol 187, 2858–2869.

Bass, K.A., and Hecht, D.W. (2002). Isolation and characterization of cLV25, a Bacteroides fragilis chromosomal transfer factor resembling multiple Bacteroides sp. mobilizable transposons. J Bacteriol 184, 1895–1904.

Brito, I.L., Yilmaz, S., Huang, K., Xu, L., Jupiter, S.D., Jenkins, A.P., Naisilisili, W., Tamminen, M., Smillie, C.S., Wortman, J.R., et al. (2016). Mobile genes in the human microbiome are structured from global to individual scales. Nature 535, 435–439.

Browne, H.P., Forster, S.C., Anonye, B.O., Kumar, N., Neville, B.A., Stares, M.D., Goulding, D., and Lawley, T.D. (2016). Culturing of ‘unculturable’ human microbiota reveals novel taxa and extensive sporulation. Nature 533, 543–546.

Chatzidaki-Livanis, M., Geva-Zatorsky, N., and Comstock, L.E. (2016). Bacteroides fragilis type VI secretion systems use novel effector and immunity proteins to antagonize human gut Bacteroidales species. Proc Natl Acad Sci U S A 113, 3627–3632.

Chin, C.S., Peluso, P., Sedlazeck, F.J., Nattestad, M., Concepcion, G.T., Clum, A., Dunn, C., O’Malley, R., Figueroa-Balderas, R., Morales-Cruz, A., et al. (2016). Phased diploid genome assembly with single-molecule real-time sequencing. Nat Methods 13, 1050–1054.

Costea, P.I., Hildebrand, F., Arumugam, M., Backhed, F., Blaser, M.J., Bushman, F.D., de Vos, W.M., Ehrlich, S.D., Fraser, C.M., Hattori, M., et al. (2018). Enterotypes in the landscape of gut microbial community composition. Nat Microbiol 3, 8–16.

Coyne, M., Roelofs, KG, Comstock, LE (2016). Type VI secretion systems of human gut Bacteroidales segregate into three genetic architectures, two of which are contained on mobile genetic elements. BMC Genomics 17.

Coyne, M., Zitomersky, N., McGuire, A., Earl, A., and Comstock, L. (2014). Evidence of extensive DNA transfer between Bacteroidales species within the human gut. mBio 5.

Fu, L., Niu, B., Zhu, Z., Wu, S., and Li, W. (2012). CD-HIT: accelerated for clustering the next-generation sequencing data. Bioinformatics 28, 3150–3152.

Garcia-Bayona, L., and Comstock, L.E. (2018). Bacterial antagonism in host-associated microbial communities. Science 361.

Groussin, M.e.a. (2020). Industrialization is associated with elevated rates of horizontal gene transfer in the human microbiome. BioRxiv.

Hecht, A.L., Casterline, B.W., Earley, Z.M., Goo, Y.A., Goodlett, D.R., and Bubeck Wardenburg, J. (2016). Strain competition restricts colonization of an enteric pathogen and prevents colitis. EMBO Rep 17, 1281–1291.

Hehemann, J.H., Kelly, A.G., Pudlo, N.A., Martens, E.C., and Boraston, A.B. (2012). Bacteria of the human gut microbiome catabolize red seaweed glycans with carbohydrate-active enzyme updates from extrinsic microbes. Proc Natl Acad Sci U S A 109, 19786–19791.

Hyatt, D., Chen, G.L., Locascio, P.F., Land, M.L., Larimer, F.W., and Hauser, L.J. (2010). Prodigal: prokaryotic gene recognition and translation initiation site identification. BMC Bioinformatics 11.

Jiang, X., Hall, A.B., Xavier, R.J., and Alm, E.J. (2019). Comprehensive analysis of chromosomal mobile genetic elements in the gut microbiome reveals phylum-level niche-adaptive gene pools. PloS one 14, e0223680.

Kolmogorov, M., Armstrong, J., Raney, B.J., Streeter, I., Dunn, M., Yang, F., Odom, D., Flicek, P., Keane, T.M., Thybert, D., et al. (2018). Chromosome assembly of large and complex genomes using multiple references. Genome Res 28, 1720–1732.

Kolmogorov, M., Yuan, J., Lin, Y., and Pevzner, P.A. (2019). Assembly of long, error-prone reads using repeat graphs. Nat Biotechnol 37, 540–546.

Koren, S., Walenz, B.P., Berlin, K., Miller, J.R., Bergman, N.H., and Phillippy, A.M. (2017). Canu: scalable and accurate long-read assembly via adaptive k-mer weighting and repeat separation. Genome Res 27, 722–736.

Krawczyk, P.S., Lipinski, L., and Dziembowski, A. (2018). PlasFlow: predicting plasmid sequences in metagenomic data using genome signatures. Nucleic Acids Res 46, e35.

Kumar, S., Stecher, G., Li, M., Knyaz, C., and Tamura, K. (2018). MEGA X: Molecular evolutionary genetics analysis across computing platforms. Mol Biol Evol 35, 1547–1549.

Marasini, D., Karki, A.B., Bryant, J.M., Sheaff, R.J., and Fakhr, M.K. (2020). Molecular characterization of megaplasmids encoding the type VI secretion system in Campylobacter jejuni isolated from chicken livers and gizzards. Scientific Rep 10, 12514.

Ofir, G., Melamed, S., Sberro, H., Mukamel, Z., Silverman, S., Yaakov, G., Doron, S., and Sorek, R. (2018). DISARM is a widespread bacterial defence system with broad anti-phage activities. Nat Microbiol 3, 90–98.

Pantosti, A., Tzianabos, A.O., Onderdonk, A.B., and Kasper, D.L. (1991). Immunochemical characterization of two surface polysaccharides of Bacteroides fragilis. Infect Immun 59, 2075–2082.

Poyet, M., Groussin, M., Gibbons, S.M., Avila-Pacheco, J., Jiang, X., Kearney, S.M., Perrotta, A.R., Berdy, B., Zhao, S., Lieberman, T.D., et al. (2019). A library of human gut bacterial isolates paired with longitudinal multiomics data enables mechanistic microbiome research. Nat Med 25, 1442–1452.

Ross, B.D., Verster, A.J., Radey, M.C., Schmidtke, D.T., Pope, C.E., Hoffman, L.R., Hajjar, A.M., Peterson, S.B., Borenstein, E., and Mougous, J.D. (2019). Human gut bacteria contain acquired interbacterial defence systems. Nature 575, 224–228.

Santoriello, F.J., Michel, L., Unterweger, D., and Pukatzki, S. (2020). Pandemic Vibrio cholerae shuts down site-specific recombination to retain an interbacterial defence mechanism. Nat Comm 11, 6246.

Schwarz, F.W., Toth, J., van Aelst, K., Cui, G., Clausing, S., Szczelkun, M.D., and Seidel, R. (2013). The helicase-like domains of type III restriction enzymes trigger long-range diffusion along DNA. Science 340, 353–356.

Seemann, T. (2014). Prokka: rapid prokaryotic genome annotation. Bioinformatics 30, 2068–2069.

Sievers, F., Wilm, A., Dineen, D., Gibson, T.J., Karplus, K., and Li, W. (2011). Fast, scalable generation of high-quality protein multiple sequence alignments using Clustal Omega. Mol Sys Biol 7.

Stamatakis, A. (2014). RAxML version 8: a tool for phylogenetic analysis and post–analysis of large phylogenies. Bioinformatics 30, 1312–1313.

Verster, A.J., Ross, B.D., Radey, M.C., Bao, Y., Goodman, A.L., Mougous, J.D., and Borenstein, E. (2017). The landscape of type VI secretion across human gut microbiomes reveals its role in community composition. Cell Host Microbe 22, 411–419 e414.

Wang, G.R., Shoemaker, N.B., Jeters, R.T., and Salyers, A.A. (2011). CTn12256, a chimeric Bacteroides conjugative transposon that consists of two independently active mobile elements. Plasmid 66, 93–105.

Waters, J.L., and Salyers, A.A. (2013). Regulation of CTnDOT conjugative transfer is a complex and highly coordinated series of events. mBio 4, e00569–00513.

Wesslund, N.A., Wang, G.R., Song, B., Shoemaker, N.B., and Salyers, A.A. (2007). Integration and excision of a newly discovered bacteroides conjugative transposon, CTnBST. J Bacteriol 189, 1072–1082.

Wexler, A.G., Bao, Y., Whitney, J.C., Bobay, L.M., Xavier, J.B., Schofield, W.B., Barry, N.A., Russell, A.B., Tran, B.Q., Goo, Y.A., et al. (2016). Human symbionts inject and neutralize antibacterial toxins to persist in the gut. Proc Natl Acad Sci U S A 113, 3639–3644.

Wick, R.R., Schultz, M.B., Zobel, J., and Holt, K.E. (2015). Bandage: interactive visualization of de novo genome assemblies. Bioinformatics 31, 3350–3352.

Wood, M.M., and Gardner, J.F. (2015). The Integration and excision of CTnDOT. Microbiol Spec 3, MDNA3-0020-2014.

Zitomersky, N.L., Coyne, M.J., and Comstock, L.E. (2011a). Longitudinal analysis of the prevalence, maintenance, and IgA response to species of the order Bacteroidales in the human gut. Infect Immun 79, 2012–2020.

Zitomersky, N.L., Coyne, M.J., and Comstock, L.E. (2011b). Longitudinal analysis of the prevalence, maintenance, and IgA response to species of the order Bacteroidales in the human gut. Infect Immun 79.

Zou, Y., Xue, W., Luo, G., Deng, Z., Qin, P., Guo, R., Sun, H., Xia, Y., Liang, S., Dai, Y., et al. (2019). 1,520 reference genomes from cultivated human gut bacteria enable functional microbiome analyses. Nat Biotechnol 37, 179–185.

